# Altitude and life-history shape the evolution of *Heliconius* wings

**DOI:** 10.1101/690396

**Authors:** Gabriela Montejo-Kovacevich, Jennifer E. Smith, Joana I. Meier, Caroline N. Bacquet, Eva Whiltshire-Romero, Nicola J. Nadeau, Chris D. Jiggins

## Abstract

Phenotypic divergence between closely related species has long interested biologists. Taxa that inhabit a range of environments and have known and diverse, natural histories, can help understand how different selection pressures shape diverging traits. In butterflies, wing colour patterns have been extensively studied, whereas wing shape diversity is less well understood despite its importance for flight. Here we study a measure of wing shape, aspect ratio, and wing size in a large dataset of over 3200 individuals, representing 13 *Heliconius* species, from across the Neotropics. We assess the relative importance of phylogenetic relatedness, natural history and habitat in determining wing shape and size. We find that both larval and adult behavioural ecology affect patterns of adult size dimorphisms. On one hand, species with solitary larvae have larger adult males, in contrast to gregarious *Heliconius* species, and indeed most Lepidoptera. On the other hand, species in the pupal-mating clade are smaller overall than those in the adult-mating clade. Interestingly, while controlling for phylogeny, sex ratios and allometry, we find that species inhabiting higher altitudes have rounder wings and, in one of the two major *Heliconius* clades, are also bigger in size than their lowland relatives. Thus, we reveal novel adaptive wing morphological divergence among *Heliconius* species beyond that imposed by natural selection on aposematic wing colouration. Our study highlights the value of phylogenetic comparative methods in study systems that have diverse and well-studied natural histories to disentangle the selection pressures shaping adaptive phenotypes.

## Introduction

Identifying the selective forces driving phenotypic divergence among closely related species lies at the core of evolutionary biology research. Adaptive radiations, in which descendants from a common ancestor rapidly fill a variety of niches, are ideal systems to investigate morphological divergence. The study of adaptive radiations has revealed that evolution often comes up with similar solutions for similar problems (Losos 2010; Marques et al. 2019). Speciose groups that have repeatedly and independently evolved convergent adaptations to life-history strategies and environments are strong systems to study selection drivers (Schluter 2000). Nevertheless, adaptive phenotypic divergence is often complex and multifaceted; with more than a single selective force in action (Maia et al. 2016; Nosil et al. 2018). For example in birds, sex differences in plumage colouration are driven by intra-specific sexual selection, while natural selection drives sexes towards more similar colourations (Dunn et al. 2015). Integrative approaches that make use of tractable traits across well-resolved phylogenies are needed to explore the selective forces driving phenotypic evolution.

Wing colouration has been the focus of considerable research effort and major strides have been made towards understanding how and when evolution leads to complex wing colour patterns, conferring aposematism, camouflage, or a mating advantage (Merrill et al. 2012; Chazot et al. 2016; Nadeau et al. 2016). Indeed, interest in wing patterns as models of evolutionary process dates back to Henry W. Bates who, in his book *The Naturalist on the River Amazon*, reflected on the astonishing diversity of butterfly wings he had observed: “It may be said, therefore, on these expanded membranes nature writes, as on a tablet, the story of the modifications of species” (Bates 1863). The dazzling diversity of colour patterns among species has perhaps obscured the less conspicuous phenotypic diversity of wing shapes and sizes (Le Roy et al. 2019), which are more often regarded as the result of sexual selection, flight trade-offs or developmental constraints (Singer 1982; Allen et al. 2011), rather than drivers of local adaptation and species diversification (but see Srygley 2004; Cespedes et al. 2015; Chazot et al. 2016).

Differences in behaviour between sexes have been identified as one of the main drivers of wing shape and size sexual dimorphism in insects (Rossato et al. 2018b; Le Roy et al. 2019). In butterflies, males tend to spend more time looking for mates and patrolling territories, while females focus their energy on searching for suitable host plants for oviposition (Rossato et al. 2018b). The same wing trait can be associated with different life history traits in each sex, resulting in sex-specific selection pressures. For example, in the Nearctic butterfly *Melitaea cinxia*, wing shape only correlates with dispersal in females, as males experience additional selection pressures that counteract selection for dispersal wing phenotypes (Breuker et al. 2007). Sex-specific behaviours can impact both wing shape and size, but differences in ecology and natural history, even across closely related species, could also have large impacts on the strength and direction of these effects (Cespedes et al. 2015; Chazot et al. 2016).

Another important source of phenotypic variation in insect wings is the abiotic environment they inhabit. Air pressure decreases with altitude, which in turn reduces lift forces required for flight. To compensate for this, insects may increase wing area relative to body size to reduce the velocity necessary to sustain flight (Dudley 2002; Dillon et al. 2018). Wing shape in the widespread *Drosophila melanogaster* has been observed to vary adaptively across latitudes and altitudes, with wings getting rounder and larger in montane habitats, possibly to, maintain flight function in lower air pressures (Stalker and Carson 1948; Pitchers et al. 2012; Klepsatel et al. 2014). In butterflies, high aspect ratios, i.e. long and narrow wings, reduce drag caused by wing tip vortices, thus lowering the energy required for flight and promoting gliding for longer distances (Le Roy et al. 2019). Variation in wing phenotypes has been detected at the microhabitat level, for example *Morpho* butterfly clades that dwell in the understory have rounder wings than canopy-specialist clades, allowing for increased manoeuvrability (Chazot et al. 2016). An extreme case of environmental effects on wing shape can be found in Lepidoptera inhabiting the windy, barren highlands of the Andes, where an interaction between behavioural sex differences and extreme climatic conditions have led to flightlessness in females of several species (Pyrcz et al. 2004).

*Heliconius* or “longwing” butterflies are an excellent example of colour pattern evolution for aposematic mimicry, where co-occurring subspecies share warning colour patterns to avoid predation, creating multi-species mimicry rings across South America (Merrill et al., 2015). Wing shape and size are part of the mimetic signal (Jones et al. 2013; Mérot et al. 2016; Rossato et al. 2018a). Wing morphology is involved in many aspects of *Heliconius* biology other than mimicry, such as mating or flight mode, but these have been considerably less well studied (but see Rodrigues and Moreira 2004; Srygley 2004; Mendoza-Cuenca and Macías-Ordóñez 2010).

The wide range of environments that *Heliconius* species inhabit, together with their diverse natural history and well-resolved phylogeny make them a good study system for teasing apart the selective forces driving wing phenotypic divergence. For example, larval gregariousness has evolved independently three times across the phylogeny, with some species laying clutches of up to 200 eggs, while the others lay eggs singly and larvae are often cannibalistic (Beltrán et al. 2007; Jiggins 2016). In gregarious *Heliconius* species, increased female size might be beneficial as the enlarged egg load might necessitate larger wings. Another striking life history trait, pupal-mating, has evolved only once, in one of the two major clades (hereafter the “erato clade”), following the most basal split in the *Heliconius* phylogeny. This mating strategy involves males copulating with females as they emerge from the pupal case, or even before (Deinert et al. 1994; Beltrán et al. 2007). Pupal-mating leads to a whole suite of distinct selection pressures but these are hard to tease apart from the effects of phylogeny due to its single origin (Beltrán et al. 2007; Jiggins 2016). Further ecological differences could arise from relatively high-altitude environmental specialisation of some *Heliconius* species, such as *H. telesiphe* and *H. hierax*, which are only found above 1000m. This is in contrast to wide-ranging species such as *H. melpomene* and *H. erato*, which can be found from 0 to 1800 m above sea-level (Rosser et al. 2015; Jiggins 2016). Potential adaptations to altitude are yet to be explored.

Here we examine variation in wing shape and size across 13 species that span most of the geographical range of the *Heliconius* genus. First, we photographed thousands of wings collected by many *Heliconius* researchers since the 1990s from wild populations across South and Central America, covering a 2100 m elevation range (Fig. 1 A). Wing dimensions for 3242 individuals, obtained from standardised images using an automated pipeline, were then used to address the following questions. (1) Are there size and shape sexual dimorphisms, and if so, do they correlate with known life-history traits? (2) To what extent are species’ wing shape and size variation explained by shared ancestry? (3) Are wing shape and size affected by the elevations species inhabit?

**Figure 1.**
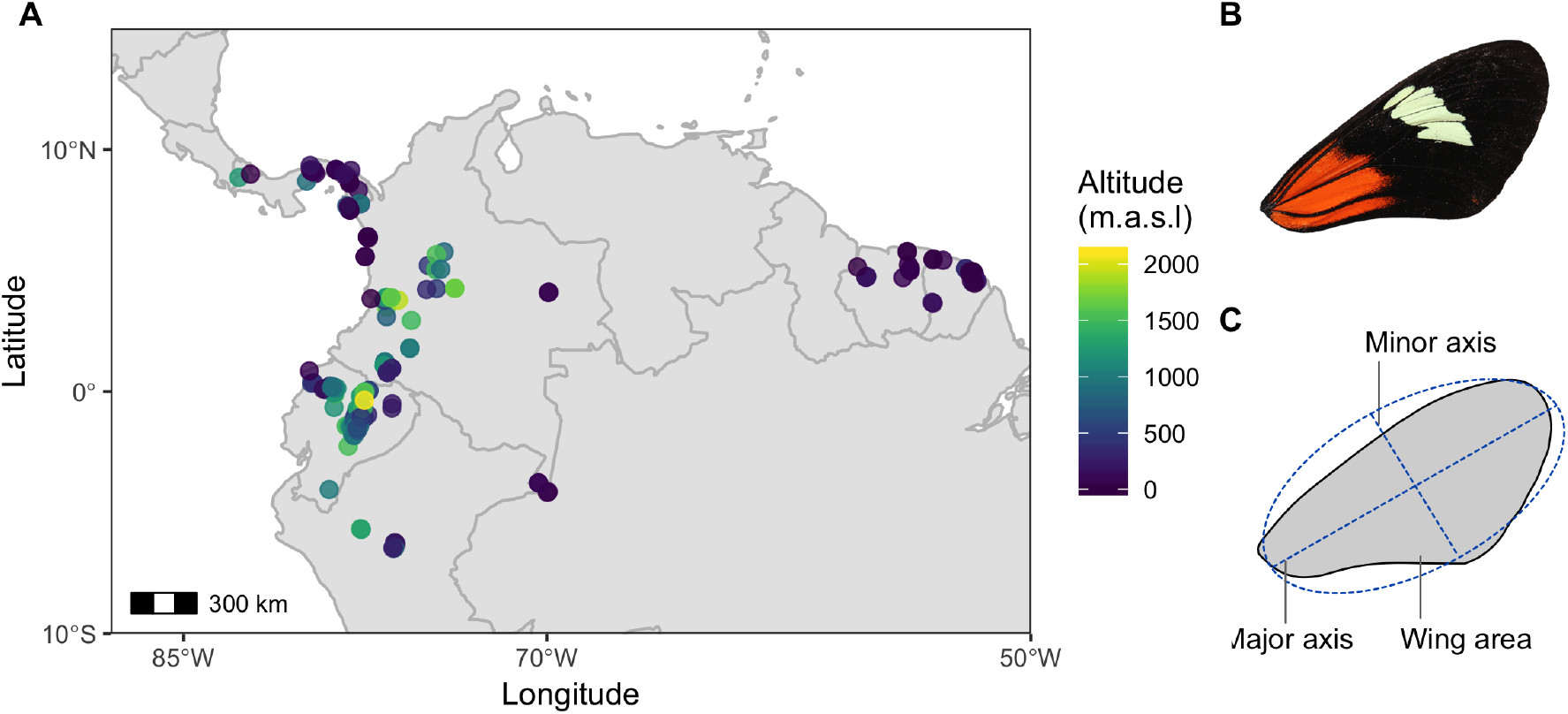
Localities and forewing measurements. (A) Map of exact locations (n=313) across South America from where the samples used for our analyses were collected. Points are coloured by altitude. (B) Representative of a right forewing image of *H. melpomene malleti.* (C) Measurements taken from each wing by fitting an ellipse with Fiji custom scripts.

## Methods

### STUDY COLLECTION

The wild specimens studied here were collected using hand nets between 1998 and 2018 in 313 localities across Panama, Colombia, Ecuador, French Guiana, Suriname, and Peru (Fig. 1 A), and stored in the Department of Zoology, University of Cambridge (Earthcape database). Collection altitudes ranged from sea level to 2100m above sea level (Fig 1 A). Detached wings were photographed dorsally and ventrally with a DSLR camera with a 100 mm macro lens in standardised conditions. All the images are available in the public repository Zenodo (https://zenodo.org/communities/butterfly/) and full records with data are stored in the EarthCape database (https://heliconius.ecdb.io).

### WING MEASUREMENTS

Damage to wings was manually scored in all the images and damaged specimens were excluded from our analyses. To obtain wing measurements from the images, we developed custom scripts for Fiji (Schindelin et al., 2012), to automatically crop, extract the right or left forewing, and perform particle size analysis (Fig. 1 B). Butterflies predominantly use their forewings for flight (Wootton 2002; Le Roy et al. 2019) and forewing and hindwing sizes are tightly correlated in this genus (Strauss, 1990), thus we only include forewings here. For wing size, we obtained total wing area (in mm^2^, hereafter “size”).

For examining wing shape, the custom scripts first fitted an ellipse to the forewings and measured the length of the longest axis and the length of the axis at 90 degrees to the former (Fig. 1 C). Aspect ratio corresponds to the length of the major axis divided by the length of the minor axis, hereafter “shape” (Fig. 1 C). The data were checked for visual outliers on scatter-plots, which were examined, and removed from the analyses if the wing extraction pipeline had failed.

### STATISTICAL ANALYSESs

All analyses were run in R V2.13 (R Development Core Team 2011) and graphics were generated with the package *ggplot2* (Ginestet 2011). Packages are specified below. All R scripts can be found in the public repository Zenodo (Zenodo: tbc), including custom Fiji scripts for wing image analysis.

#### Sexual dimorphism

Species and sexes mean trait values were calculated for the 13 *Heliconius* species in our study that had more than 30 individuals each and for which accurate locality and altitude data were available (SI: Table 1, 16 species were excluded), resulting in a dataset of 3243 individuals. Sexual dimorphism in size and shape was estimated as the female increase in mean trait values with respect to males, thus negative values represent larger trait values in males, and vice-versa.

**Table 1.** Summary of model outputs derived from the analyses of size sexual dimorphism (positive values represent species with bigger females, negative represent species with bigger males), and shape and size phylogenetic generalised least squares. For the latter, summary statistics are presented for the overall model and the explanatory variable of interest, altitude. Full model tables for PGLS can be found in the Supplementary Materials (Table S2).

| Response variable (wing trait) | Trait repeat-ability (R) | Model type | Corr. structure | Fixed effects | Est. | S.E. | t-value | predictor p-value | d.f. (res. d.f.) | Adj. R<sup>2</sup> |
| --- | --- | --- | --- | --- | --- | --- | --- | --- | --- | --- |
| Sexual size dimorphism | NA | lm | Gaussian | (Intercept)<br>Solitary larvae | 5.12<br>-10.3 | 1.28<br>1.74 | 4.00<br>-5.9 | 0.002***<br>0.0001*** | 1<br>(11) | 0.76 |
| Shape | 0.74<br>(p=0) | PGLS<br>(nmle) | Phylogeny,<br>intra-sp<br>variance,<br>sample size | (Intercept)<br>Altitude | 0.15<br>-2.2E-4 | 0.53<br>7.9E-5 | 0.27<br>-2.75 | 0.79<br>0.028* | 13<br>(7) | NA |
| Size | 0.47<br>(p=0) | PGLS<br>(nmle) | Phylogeny,<br>intra-sp<br>variance,<br>sample size | (Intercept)<br>Altitude | -705.4<br>1.3 | 392.1<br>0.49 | -1.79<br>2.63 | 0.13<br>0.047* | 13<br>(5) | NA |

We modelled variation in size and shape sexual dimorphism with ordinary least squares (OLS) linear regressions, implemented in the ‘lm’ function. For both models, predictor variables initially included larval gregariousness of the species (following Beltrán et al.

2007), mating strategy (pupal-mating vs. adult-mating clade), species mean shape and size, species sex ratio in our dataset, and species sexual wing shape or size dimorphism (respectively). We used backward selection to remove terms according to level of significance, retaining terms at a threshold of α = 0.05. We report the overall variation explained by the fitted models (R^2^) and the relative contributions of each explanatory variable (partial R^2^) (Grömping 2006).

#### Phylogenetic signal

We calculated the phylogenetic signal index Abouheif’s Cmean (Abouheif 1999) for mean wing shape, size and species altitude, which is an autocorrelation metric suitable for datasets with a relatively low number of species and that does not infer an underlying evolutionary model (Münkemüller et al. 2012). Observed and expected distribution plots for phylogenetic signal estimates are shown in the Supplementary Materials and were computed with the package *adephylo* (Jombart and Dray 2010). We used a pruned tree with the 13 species under study from the most recent molecular *Heliconius* phylogeny (Kozak et al., 2015). We plotted centred trait means across the phylogeny with the function barplot.phylo4d() from the package *phylosignal* (Keck et al. 2016). To test and visualise phylogenetic signal further, we built phylocorrelograms for each trait with the function phyloCorrelogram() of the same package, which estimates Moran’s I autocorrelation across matrices with varying phylogenetic weights. Then, the degree of correlation (Morans’ I) in species trait values can be assessed as phylogenetic distance increases (Keck et al. 2016).

#### Variation across species

Species wing trait means may be correlated due to shared ancestry (Freckleton et al. 2002; Chazot et al. 2016). Therefore, to explore the effects of the environment on the traits under study, models that incorporate expected correlation between species are required. One of the most commonly employed and versatile techniques to do this is phylogenetic generalised least squares (PGLS). Although often ignored, these models assume the presence of phylogenetic signal on the model residuals of the trait under study (here wing shape or size) controlling for all potential covariates, and not just phylogenetic signal on the species mean trait values (Revell 2010; Garamszegi 2014). Thus, we estimated phylogenetic signal as described above (Keck et al. 2016) with the residuals of a generalised least squares (GLS) model that included wing shape or size as response variables and potential allometric, environmental and natural history terms as explanatory variables. We obtained phylogenetic correlograms for trait residuals (presented in the SOM), and centred trait residuals for plotting across the phylogeny as detailed above for trait means (Keck et al. 2016).

Phylogenetic comparative methods assume that species-specific mean trait values are a good representation of the true trait values of the species under study, in other words, that the within-species variation is negligible compared to the across-species variation (Garamszegi 2014). To test this, we first used an ANOVA approach, with species as a factor explaining the variation of mean trait values. We then estimated within-species trait repeatability, or intra-class correlation coefficient (ICC), with a linear mixed model approach that had species as a random effect, specifying a Gaussian distribution and 1000 parametric bootstraps to quantify uncertainty, implemented with the function rptGaussian() in *rptR* package (Stoffel et al. 2017). By specifying species as a random effect, the latter approach estimates the proportion of total trait variance accounted for by differences between species. A trait with high repeatability indicates that species-specific trait means are reliable estimates for further analyses (Stoffel et al. 2017). We, nevertheless, accounted for within-species variation in the models described below.

To examine the effects of altitude on wing shape and size variation between species, we first tested for phylogenetic signal in these traits. Significant phylogenetic signal was detected in the residuals of wing shape and size regression models (SOM Fig. S1, Fig. S2), so we used maximum log-likelihood PGLS regression models with the trait phylogenetic correlation fitted as a correlation term, implemented with the gls() function from the *nmle* package (Pinheiro et al. 2007). We assumed a Brownian motion model of trait evolution for both traits, by which variation across species accumulated along all the branches at a rate proportional to the length of the branches (Freckleton et al. 2002). To select the most supported model given the available data, i.e. one that improves model fit while penalising complexity, we used the Akaike information criterion (AIC); (Anderson, 2008; Garamszegi, 2014), where the best models had the lowest AIC values. Thus, to select covariates to include in the model, we used step backward and forward selection based on AIC with the function stepAIC(), from the *MASS* package (Ripley, 2011; Zhang, 2016). Maximal PGLS models included species mean altitude and distance from the Equator, sex ratio in our samples interacting with sexual size dimorphism, as well as either wing shape or size to control for potential allometric relationships, which could be different among closely-related taxa (Outomuro and Johansson 2017). Interaction terms between altitude and size/shape were also included in the maximal models when relevant. Minimal PGLS models consisted of the trait under study explained solely by its intercept, without any fixed effects. We weighted PGLS regressions to account for unequal trait variances and sample sizes across species (for sample sizes and standard errors of species’ trait means see SOM Table 1). This was achieved by modifying the correlation structure of the model with combined variances obtained with the function varFixed() and specified with the argument “weights” (Pinheiro et al. 2007; Paradis 2012; Garamszegi 2014).

We examined the interactions between fixed effects and phylogenetic correlations and their explanatory power in the selected models by deriving partial R^2^ based on maximum likelihood estimates from full and reduced models, implemented in R with:

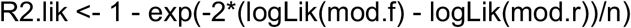

where mod.f is the full model, mod.r is the reduced model with either fixed effects removed or phylogenetic correlation removed, and n is the species sample size (Paradis 2012; Ives 2018). To visualise the effects of altitude on wing shape and size we plotted the residuals of a PGLS model built as described above without altitude as an explanatory variable, against the species mean altitude. Additionally, to understand the effect of controlling for relevant fixed effects, phylogenetic correlation, intraspecific variance, and sample size, we present the raw wing shape and size species variation across altitudes in the Supplementary Materials.

## Results

We obtained intact-wing measurements for 3243 individuals of 13 *Heliconius* species from across the phylogeny and from over 300 localities (Fig. 1, Table S1). We have made all of these wing images publicly available at the Zenodo repository.

### SEXUAL DIMORPHISM

Sexual dimorphism in wing size was found throughout the phylogeny, but in opposing directions in different species (Fig. 2). Mean sizes were significantly different (P<0.05) among sexes in all of the six best sampled species (see Table S1 for t-test summary statistics), indicating that the marginally- or non-significant trends in other species probably reflect a lack of power caused by low numbers of females. The six species with trends toward larger females have gregarious larvae (pink, Fig. 2), whereas the six species with trends toward larger males lay eggs singly (black, Fig. 2). Larval gregariousness alone explained 66.5% of the total natural variation in sexual size dimorphism across species (Table 1; Gaussian LM: F_1,11_= 24.9, P<0.001, R^2^=0.67). There was a significant phylogenetic signal in sexual size dimorphism (Abouheif’s Cmean=0.28, P=0.03; SI, Fig. S1). This would be expected from the evolutionary history of gregariousness, as it has evolved independently at least three times across the *Heliconius* phylogeny (Beltrán et al. 2007). However, even when accounting for phylogenetic correlation in the model, larval gregariousness remained a significant predictor of size sexual dimorphism (SI, Table S2).

**Figure 2.**
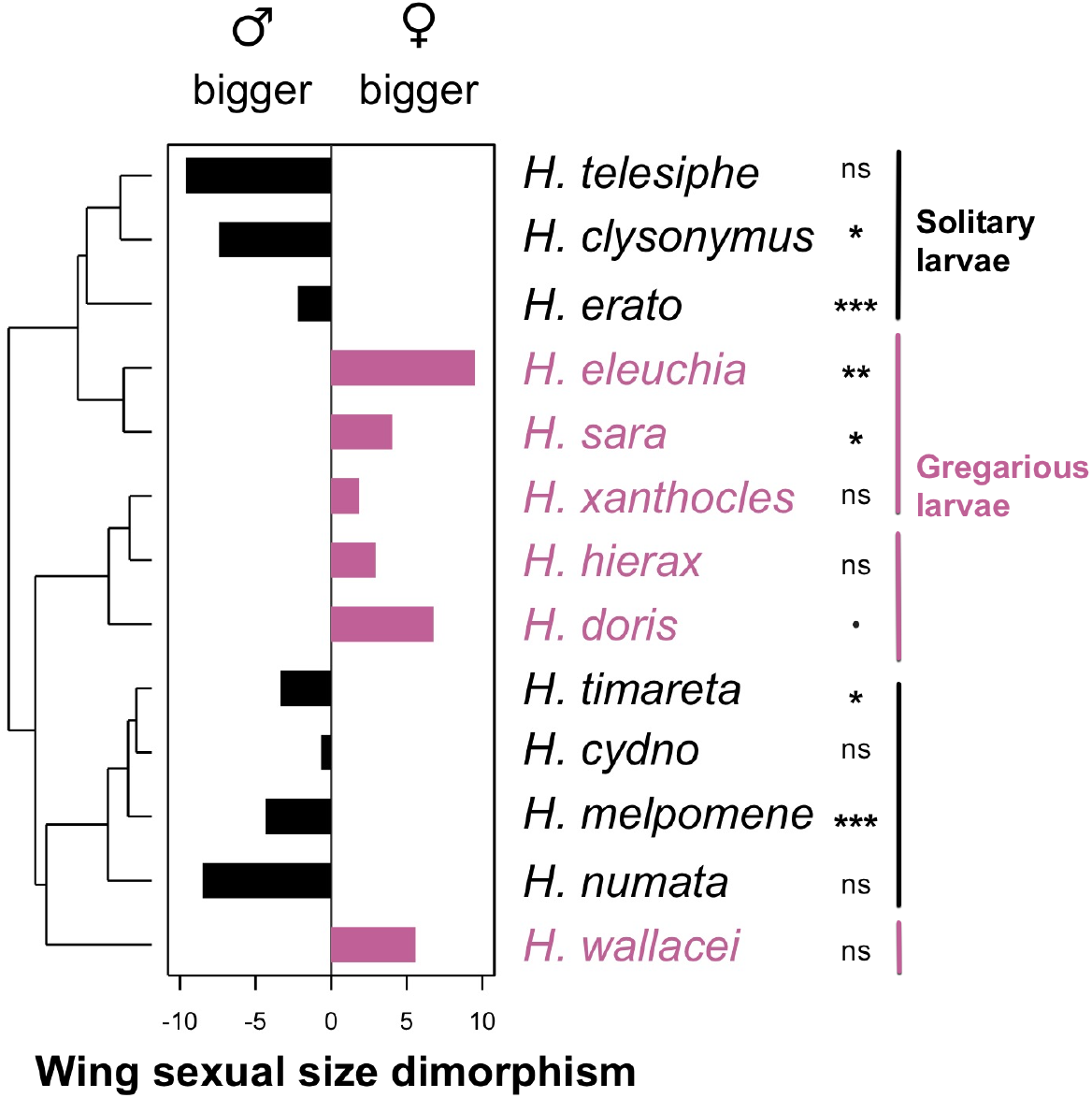
Wing size sexual dimorphism variation across the phylogeny. Wing size was measured as total right or left forewing area (in mm^2^). Bar plot represents sexual size dimorphism calculated as percentage difference in female vs. male size (positive means bigger females). Species with gregarious larvae are coloured in pink, and those with solitary larvae are coloured in black. Stars represent significance levels of two sample t-tests between female and male wing sizes for each species (•<0.1, *< 0.05, **<0.01, ***<0.001), for full t-tests output see Table S1.

Sexual dimorphism in wing shape was found only in two species from the erato clade, the widespread *H. erato* had longer-winged males whereas the high-altitude specialist *H. eleuchia* had longer-winged females (T-test, *H. erato:* t_739_=-9.52, P<0.0001, *H. eleuchia:* t_49_=2.21, p<0.05). Wing shape sexual dimorphism across species could not be explained with the variables here studied and had no phylogenetic signal (Abouheif’s Cmean=0.02, P=0.2; SI, Fig. S1).

### PHYLOGENETIC SIGNAL

The 13 *Heliconius* species studied differed significantly in wing size and shape (ANOVA, Shape: F_12, 3230_ = 215.8, P < 0.0001, Size: F_12, 3230_ = 189.3, P < 0.0001). We estimated within-species trait repeatability to assess their reliability as species mean estimates for phylogenetic analyses. Wing shape had higher intra-class repeatability than wing size, with 74% and 47% of the total shape and size variance explained by differences between species, respectively (Shape: R=0.74, S.E.=0.09, P<0.0001; Size: R=0.47, S.E.=0.1, P<0.0001). We estimated intra-class repeatability for males and females separately to remove the potential effect of size sexual dimorphism on trait variation, and male size repeatability remained much lower than male wing shape repeatability (Male shape: R=0.75, S.E.=0.08, P<0.0001; Male Size: R=0.51, S.E.=0.1, P<0.0001).

Mean wing shape showed no phylogenetic signal (Abouheif’s Cmean=0.15, P=0.1; SI: Fig. S1, Fig. S2 B), whereas mean wing size showed a strong phylogenetic signal (Fig. 3, Abouheif’s Cmean=0.33, P=0.01; SI: Fig. S1, Fig. S3 B). Wing sizes of species in the melpomene clade were on average 14.6% larger than those of species in the erato clade, with *H. timareta* being 56% larger than *H. sara* (Fig. 3 C, *timareta:* mean=605.8 mm^2^, s.e.=3.6; *sara:* mean=386 mm^2^, s.e=3.1). Nevertheless, when controlling for fixed effects on wing shape and size (species mean: size/shape, altitude, latitude, sex ratio, size sexual dimorphism), the residuals of both traits show a phylogenetic signal (SI: Fig. S2 A, C-Shape residuals: Abouheif’s Cmean=0.31, P<0.05; Fig. S3 A, C-Size: Abouheif’s Cmean=0.21, P=0.05).

**Figure 3.**
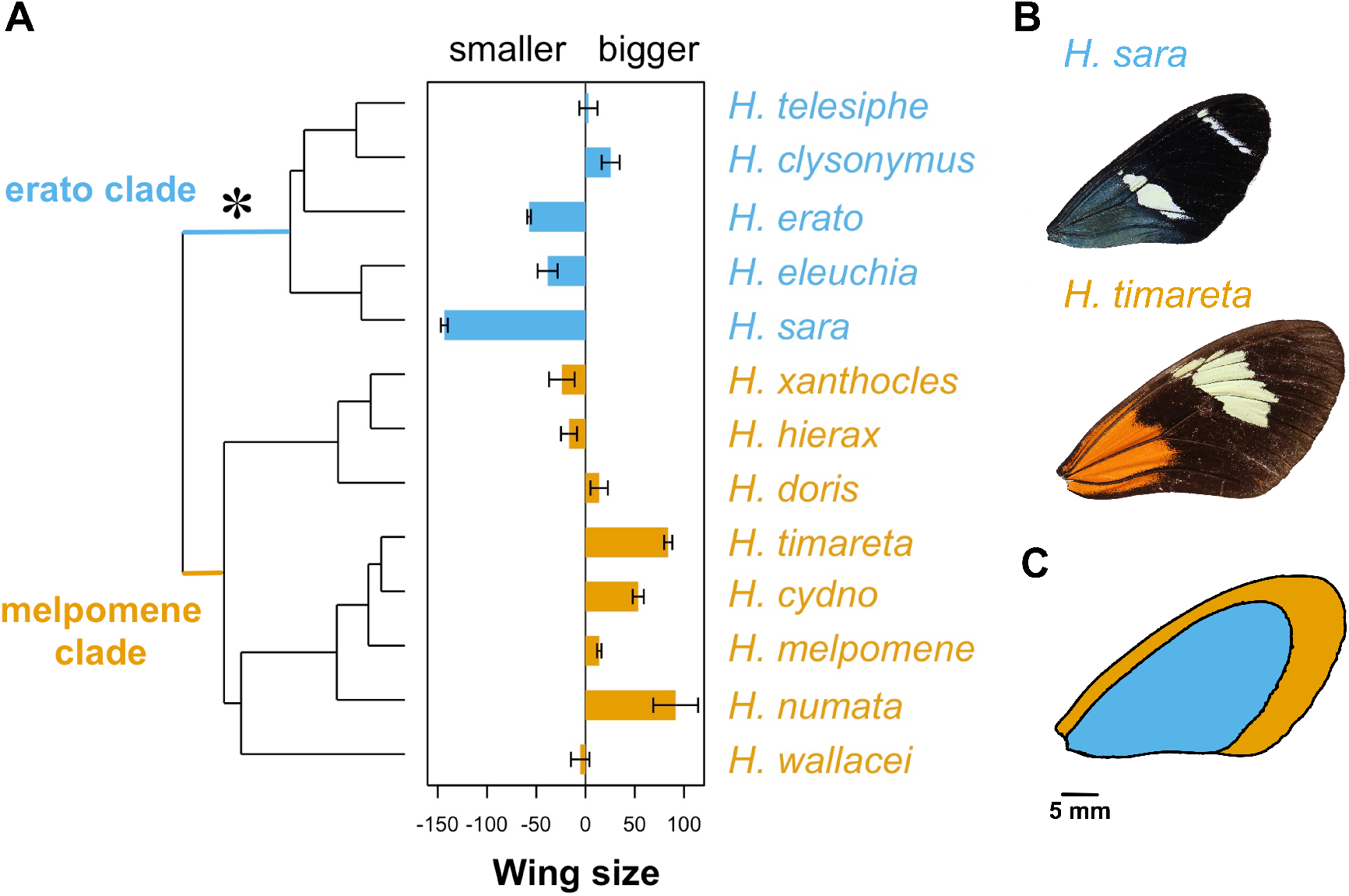
Male wing size differences across the phylogeny. (A) Bar plot represents centred mean wing size per species (positive values represent species with bigger wings than the mean *Heliconius* wing). Wing size, x-axis, is the difference in wing size from the mean (in mm^2^). Error bars represent standard errors. The star represents the origin of pupal-mating. Species from the erato clade are in blue, and those from the melpomene clade are in orange. (B) Representatives of *H. timareta* and *H. sara* closest to the mean wing size of the species are shown (606.25 mm^2^ and 386.6 mm^2^, respectively). (C) Images from (B) superimposed to compare visually the mean size difference between the two species.

### PATTERNS ACROSS ALTITUDES

To account for the moderate within-species repeatability of wing size, we included intra-specific variance in the error structure of the phylogenetic generalised least squares (PGLS) models for both wing traits. Altitude had an effect on wing size and shape (Table 1). Species wings got rounder with increasing altitudes both when accounting for fixed effects and phylogeny (Table 1, full model Table S2, Fig. 4). These trends were also evident when examining raw mean wing shapes (Fig. S3 A, Gaussian LM: F_1, 9_ = 5.37, P < 0.05, R^2^=0.30), except in the *H. telesiphe* and *H. clysonymus* highlands clade, which showed significant phylogenetic autocorrelation (Moran’s I index: *H. clysonymus* 0.53, *H. telesiphe* 0.49). Altitude had similarly strong effects on wing shape residuals for the erato and melpomene clades (Fig. 4A, Table S2), but there were further clade-specific effects, i.e. evolutionary rate shifts, that could not be accounted for by phylogeny alone in the PGLS models and that are represented by the different intercepts in Fig. 4A.

**Figure 4.**
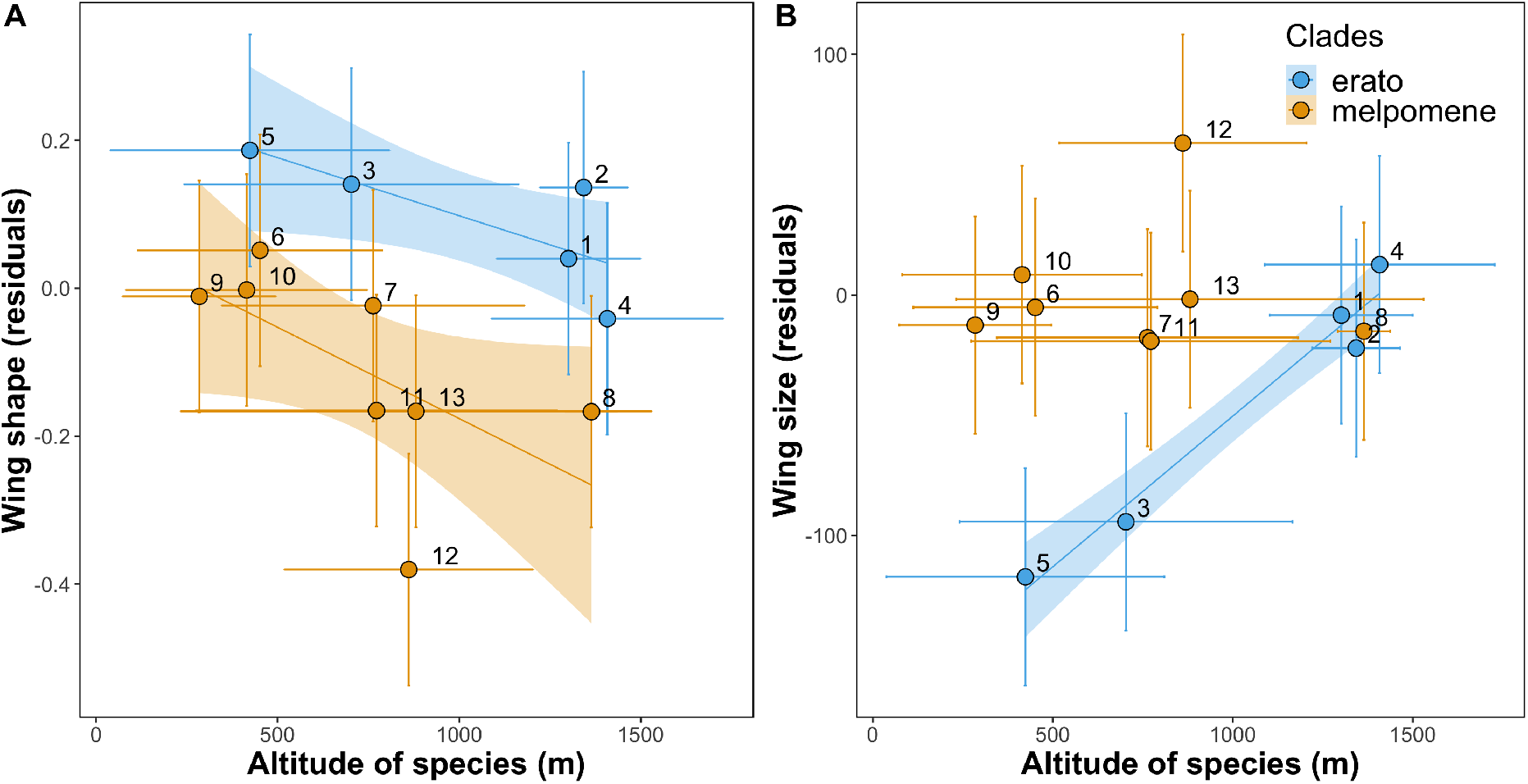
Species variation in wing shape (A) and wing size (B). Residual plots show the effect of altitude (meters above sea level) on wing shape (aspect ratio) and wing size (area mm2) residuals of PGLS model controlling for selected fixed effects (Table S2), intraspecific variance, sample size, and phylogenetic correlation. Points represent species mean values. Horizontal and vertical lines show standard error for species mean altitude and mean trait residuals, respectively. Lines show best linear fit for significant effects and are coloured by clade (blue: erato clade, orange: melpomene clade). Shaded areas show confidence bands at 1 standard error. Numbers correspond to *Heliconius* species: 1 *H. telesiphe*, 2 *H. clysonymus*, 3 *H. erato*, 4 *H. eleuchia*, 5 *H. sara*, 6 *H. doris*, 7 *H. xanthocles*, 8 *H. hierax*, 9 *H. wallacei*, 10 *H. numata*, 11 *H. melpomene*, 12 *H. timareta*, 13 *H. cydno*.

Species wing sizes increased with elevation (Table 1, full model Table S2). High altitude species of the erato clade were bigger than their lowland sister species, both with PGLS residuals (Fig. 4B, blue, Gaussian LM: F_1,10_= 118, R^2^=0.97, p=0.001) and with raw wing size (Fig. S3 B, blue, Gaussian LM: F_1,10_= 17.1, R^2^=0.80, p=0.03). When examining the melpomene clade patterns alone, we did not detect a size-altitude relationship (Fig. 4B). At the extremes of the erato clade elevational range were the highland species *H. clysonymus*, which was 39% bigger than the lowland species *H. sara* (Fig. S3 B). The phylogeny term of the wing size model interacted with the fixed effects, and improved the fit and predictive power of the model (Table S4), as expected for a trait with a strong phylogenetic signal (Revell 2010; Ives 2018).

## Discussion

The fascination of H. W. Bates for butterfly wing colouration has stimulated several generations of research and *Heliconius* wing patterns have proven to be excellent study systems for understanding evolution and speciation. Here we have extended this research by examining wing shape and size variation among more than 3200 individual butterflies, across sexes, clades, and altitudes in 13 species of *Heliconius* butterflies. We have shown that a large proportion of female biased sexual size dimorphism can be explained by the evolution of larval gregariousness, and that male biased sexual size dimorphism is present only in species that lay eggs singly, regardless of their mating strategy. For the first time in this system, we describe wing morphological variation across environmental clines, with species found at higher altitudes consistently having rounder wings. Here we demonstrate that *Heliconius* wing size and shape is affected by a plethora of behavioural and environmental selection pressures, in addition to those imposed by Müllerian mimicry.

### WING SHAPE VARIATION

Wing shape in butterflies, and other flying animals (Farney and Fleharty 1969; Buler et al. 2017), determines flight mode and speed, and is therefore predicted to vary with life-history requirements across sexes and species. Long wings are generally associated with faster gliding flying, whereas round wings with low aspect ratio values favour slow but more manoeuvrable flight motions (Betts and Wootton 1988; Chai and Srygley 1990; Chazot et al. 2016; Le Roy et al. 2019). For instance, monarch butterfly populations with longer migrations have more elongated wings than resident populations (Satterfield and Davis 2014), and males of *Morpho* species that dwell in the canopy also have higher aspect ratios to glide faster through open areas (DeVries et al. 2010). In contrast, female *Morpho* butterflies tend to have rounder wings, and shape sex differences are stronger in species with colour dimorphism, as varying crypsis may require specific flight behaviours (Chazot et al. 2016).

*Heliconius* are not notoriously sexually dimorphic, especially when compared to other butterflies such as *Morpho* (Chazot et al. 2016; Jiggins 2016). However, there are important behavioural differences between the sexes. Females are thought to have different flight habits, as they spend much of their time looking for specific host plants for oviposition (Dell’Aglio et al. 2016), or precisely laying eggs on suitable plants, while males tend to patrol open areas searching for receptive females and visit flowers more often (Joron 2005; Jiggins 2016). Thus, it might be predicted that females should have lower aspect ratios, i.e. rounder wings, than males (Jones et al. 2013). However, we only found two species, both of which belong to the pupal-mating clade, with significant, but opposing, sexually dimorphic wing shapes (males: *H. erato* longer, *H. eleuchia* rounder). This result might simply reflect strengthened intra-sexual selection in males of species that can both pupal-mate and adult-mate in the wild. In addition, the relatively low collection numbers of female *Heliconius* could hinder the detection of subtle wing shape differences across the sexes.

Sexual selection has long been known to affect wing colour pattern in *Heliconius*, as it is used for mate recognition and choice (Merrill et al. 2012). More recently, wing shape has been shown to be part of the mimetic warning signal in *Heliconius* and their co-mimics (Jones et al. 2013), as it determines flight motion and defines the overall appearance of the butterfly (Srygley 1994, 2004). For instance, wing shapes between two different morphs of *H. numata* differed consistently across their overlapping ranges, in parallel with their respective and distantly related *Melinea* co-mimics (Jones et al. 2013). However, within-morph wing shape variation was observed across the altitudinal range of *H. timareta* in Peru (Mérot et al. 2016), and in the *Heliconius* postman mimicry ring in Brazil significant across-species wing shape differences were also found (Rossato et al. 2018a). These studies highlight that while it is clear that colour pattern and, to some extent, flight are important for mimicry in *Heliconius*, wing shape is also subject to other selection pressures (Mérot et al. 2016; Rossato et al. 2018b).

We also found that species inhabiting higher altitudes have rounder wings, after accounting for phylogeny, sample size and intra-specific variance (Fig. 4 A). Rounder wings aid manoeuvrability and are associated with slower flight in butterflies (Berwaerts et al. 2002; Le Roy et al. 2019) and slower flights are generally associated with a decrease in ambient temperature (Gilchrist et al. 2000). In addition, air pressure, which directly reduces lift forces required to offset body weight during flight (Dillon 2006), decreases approximately 12% across the mean altitudinal range of the species here studied. Thus, the rounder wings in high altitude *Heliconius* species may aid flying in dense cloud forests, where increased manoeuvrability could be beneficial, or compensating for lower air pressure at higher altitude.

### WING SIZE VARIATION

Wing size showed significant sexual dimorphism in more than half of the species studied here, but some species had larger males and others larger females (Fig. 2). In most butterflies, females are larger than males, presumably because fecundity gains of increased body size are greater for females (Allen et al. 2011). Indeed, the *Heliconius* species in this study that tended to have larger-winged females were those that lay eggs in large clutches (Fig. 2), ranging from 30-200 eggs and whose larvae are highly gregarious (Beltrán et al. 2007). Thus, females of these species are likely investing more resources in fecundity than males, which leads to larger body and wing sizes that allow them to carry and lay eggs in clutches throughout adulthood. Selection for larger females is generally constrained by a trade-off between the benefits of increased fecundity at the adult stage and the higher predation risk at the larval stage associated with longer development times (Allen et al. 2011). This constraint might be alleviated in the unpalatable larvae of *Heliconius*, as bigger larval and adult size could increase the strength of the warning toxic signal to predators (Jiggins 2016).

An extensive survey identified that only six percent of lepidopteran species exhibit male-biased sexual size dimorphism, and that these patterns were generally explained by male-male competition (i.e. intrasexual selection), in which larger males had a competitive advantage (Stillwell et al. 2010; Allen et al. 2011). In contrast, nearly half of the *Heliconius* species studied here have male-biased sexual size dimorphism, and all of these lay eggs singly and have solitary larvae (Fig. 2). Male-male competition is high for *Heliconius* species, as females rarely re-mate despite their very long reproductive life-spans (Merrill et al. 2015). In addition, large reproductive investments in the form of nuptial gifts from males can, in principle, explain male-biased sexual size dimorphisms, as is the case in the polyandrous butterfly *Pieris napi* whose male spermatophore contains the amount of nitrogen equivalent to 70 eggs (Karlsson 1998; Allen et al. 2011). Male *Heliconius* spermatophores are not only nutrient-rich, but also loaded with anti-aphrodisiac pheromones that prevent re-mating of fertilised females (Schulz et al. 2008; Merrill et al. 2015). Therefore, it seems likely therefore that in species that lay eggs singly, sexual selection favouring larger males exceeds selection pressures for the large female size needed to carry multiple mature eggs. To our knowledge, *Heliconius* is the first example of a butterfly genus in which both female- and male-biased size dimorphism are found and can be explained by contrasting reproductive strategies.

We found a strong phylogenetic signal for wing size, with species from the erato clade being on average 12% smaller than those in the melpomene clade (Fig. 3). There are many ecological factors that could explain this pattern, and all could have contributing effects that are hard to disentangle (Fig. 3). Firstly, the erato clade is characterised by facultative pupal-mating (Beltrán et al. 2007; Jiggins 2016), by which males fight for pupae, guard them, and mate with females as they are emerging from the pupal case (Deinert et al. 1994; Jiggins 2016). This may increase male intra-sexual competition where smaller males that outcompete others for a spot on the female pupal case more successfully inseminate emerging females compared to larger, less agile males, and remove the potential choice of females for larger males (Deinert et al. 1994).

Secondly, pupal-mating seems to have far-reaching impacts on species life-histories (Boggs 1981). Pupal-maters are largely monandrous, with an extensive survey only finding 3 out of 251 wild *H. erato* with signs of re-mating in their reproductive tract, i.e. two or more spermatophores (Walters et al. 2015). In contrast, species in the melpomene or adult-mating clade are polyandrous, which leads to selection favouring large spermatophores (Boggs 1981), as seen in the polyandrous *Pieris napi* (Karlsson 1998). These can take up a remarkable proportion of the male abdomen, and provide mated females with abundant nutritional resources and defences that prevent them from re-mating with other males (Cardoso et al. 2009; Cardoso and Silva 2015). This could decrease selection pressure for larger males in the pupal-mating clade, as nuptial gifts need not be so large or nutrient/defence rich, leading to smaller male and female offspring. However, recent studies question the relative frequency of pupal-mating in the wild vs. adult mating encounters (Thurman et al. 2018), and others have found that the level of female emergence asynchrony determines the reproductive strategy of males (Mendoza-Cuenca and MacÍas-Ordóñez 2010). Furthermore, the single origin of pupal-mating in *Heliconius* (Fig. 2) makes it challenging to infer the impacts of this mating strategy on wing morphology, as the behaviour is confounded by phylogeny.

Finally, host-plant use varies considerably, with the erato clade generally having a more restricted and specialised diet than the melpomene clade, which feed on most lineages of *Passiflora* (Kozak et al. 2015; Jiggins 2016; de Castro et al. 2018). For example, size and female fecundity in *H. erato phyllis* has been shown to vary with host plant-use and seasonal availability across its extensive Brazilian range (Rodrigues and Moreira 2002, 2004). Thus, adult size could be largely a result of the quality host plants consumed at the larval stage.

In the erato clade larger species are found at higher altitudes (Fig. 4 B), unlike the melpomene clade species here studied. For example, the melpomene clade high-altitude specialist *H. hierax* is smaller than many of its lowland relatives and, interestingly, clusters with the other high-altitude specialists from the smaller erato clade (Fig. 4 B, species 8). Two major environmental factors are known to affect insect size across altitudinal clines. One is temperature, such that at lower temperatures, development times are longer and insects grow larger (Chown and Gaston 2010). This perhaps explains cases of Bergmann’s rule among ectotherms, where larger species are found in colder climates (Shelomi 2012; Classen et al. 2017). The generality of Bergmann’s rule for insects has been widely studied and debated, both across latitudinal and altitudinal clines, and it is highly dependent on sampling design and the level of biological organization studied (Shelomi 2012). Additionally, wing beat frequency tends to be lower at low temperatures, so larger wings are required to compensate and gain the extra lift required for flight, as seen in *Drosophila robusta* (Azevedo et al. 2006; Dillon 2006). A second factor likely to contribute to altitude related differences in wing size is air pressure changes and the correlated lower oxygen availability, which affects flight motion and kinematics as well as many physiological processes. High-altitude insects can minimise the impacts of lower air pressure by having larger wings, because this lowers the velocity required to induce flight (Dudley 2002). To assess the relative impacts of these and other environmental factors on *Heliconius* wing sizes and their relation to body sizes would require common-garden rearing and flight assays in controlled conditions.

### HERITABILITY

Our study demonstrates that multiple selective forces influence *Heliconius* wing size and shape. One important issue in phenotypically diverging populations is distinguishing between highly condition dependent or phenotypically plastic traits as opposed to heritable, conserved traits. In this study we found that 74% of the variation in wing shape could be explained by species identity, in contrast to 47% of the variation in wing size. High intra-class repeatability is often considered an indication of high trait heritability (Nakagawa and Schielzeth 2013). Additionally, wing shape is predicted to be phenotypically more constrained than size, because subtle shape variation can have large impacts on flight motion and kinematics (Jones et al. 2013; Le Roy et al. 2019).

In *Drosophila*, the genetic architecture of wing shape appears to be complex and differs (Gilchrist and Partridge 2001), and is independent of that of wing size (Carreira et al. 2011). Variability of wing size dramatically reduced when flies were reared in controlled conditions compared to wild populations, but both wing shape and size had a detectable heritable component (Bitner-Mathé and Klaczko 1999; Klaczko 1999). The patterns of variation in size across altitudes or latitudes are often not due to phenotypic plasticity, as many studies have shown their retention when populations are reared in common-garden conditions (Chown and Gaston 2010). In Monarch butterflies, for example, common-garden reared individuals from wild populations that had different migratory habits showed a strong genetic component for both wing shape and size (Altizer and Davis 2010). We have shown that different selection pressures affect wing shape and size and that the strength of these potentially varies across sexes and environmental clines (Altizer and Davis 2010).

### CONCLUSIONS

Here we have demonstrated how an understanding of natural and evolutionary history can help to disentangle the putative agents of selection on an adaptive trait. Wing trait differences across sexes, clades and environments give insight into the selective forces driving phenotypic divergence in *Heliconius*, beyond the effects of natural selection imposed by Müllerian mimicry. Our study highlights the complexity of selection pressures affecting seemingly simple traits and the need for a thorough understanding of life history differences among species.

## Acknowledgements

We would like to thank all the *Heliconius* researchers and field assistants that have contributed to the wing collection used in this study. We also thank Simon H. Martin and Stephen H. Montgomery for useful comments on early versions of this manuscript. G.M.K. was supported by a Natural Environment Research Council Doctoral Training Partnership. J.E.S. was supported by the Scurfield Memorial Bursary (University of Sheffield). Funding was provided to C.B. by the Spanish Agency for International Development Cooperation (AECID, grant number 2018SPE0000400194).

