## Supplemental Information for "Altitude and life-history shape the evolution of *Heliconius* wings"

### Supplementary Materials

**Table S1** Study species summary data. Sample sizes and wing parameters for the 13 study species, ordered phylogenetically based on the most recent *Heliconius* phylogeny (Kozak *et al.*, 2015). Size sexual dimorphism (SSD) two-sample t-tests summary statistics (last three columns). Positive t-values indicate bigger females (Fig. 2, main text).

| Species | N | Size mean ($mm^2$) | Shape mean | Altitude mean (m.a.s.l.) | Sex Ratio (%male) | SSD t-value | SSD d.f. | SSD p-value |
|---|---|---|---|---|---|---|---|---|
| *H. telesiphe* | 49 | 519.9 | 2.34 | 1307 | 88.6 | -1.40 | 5 | ns |
| *H. clysonymus* | 56 | 539.5 | 2.32 | 1348 | 69.1 | -2.05 | 25 | <0.05* |
| *H. erato* | 1576 | 465.1 | 2.09 | 824 | 72.6 | -2.83 | 748 | <0.01** |
| *H. eleuchia* | 103 | 500.0 | 2.03 | 1403 | 70.9 | 2.66 | 60 | <0.01** |
| *H. sara* | 220 | 386.6 | 2.17 | 392 | 71.8 | 2.17 | 100 | <0.05* |
| *H. xanthocles* | 35 | 502.7 | 2.02 | 995 | 69.0 | 0.35 | 13 | ns |
| *H. hierax* | 37 | 512.1 | 2.08 | 1381 | 77.8 | 0.53 | 8 | ns |
| *H. doris* | 39 | 547.1 | 2.30 | 423 | 81.1 | 1.99 | 10 | 0.07 • |
| *H. timareta* | 137 | 605.8 | 2.04 | 824 | 82.9 | -2.28 | 34 | <0.05* |
| *H. cydno* | 111 | 578.1 | 2.08 | 900 | 87.4 | -0.24 | 17 | ns |
| *H. melpomene* | 800 | 535.0 | 2.05 | 856 | 80.9 | -4.40 | 221 | <0.001*** |
| *H. numata* | 34 | 603.0 | 2.13 | 320 | 65.4 | -1.29 | 15 | ns |
| *H. wallacei* | 46 | 525.9 | 2.18 | 350 | 78.3 | 1.36 | 14 | ns |

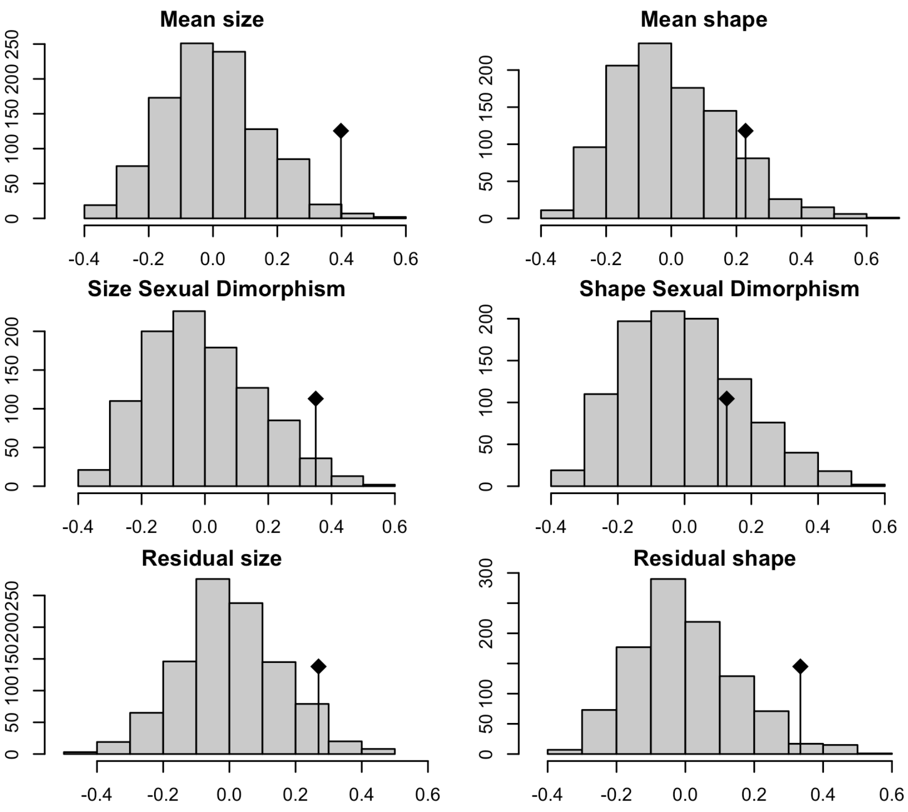


**Figure S1.** Abouheif C-mean distribution plots for six variables. Black dots depicts the observed C-mean statistic relative to the null hypothesis of randomisations along the tips of the phylogeny.

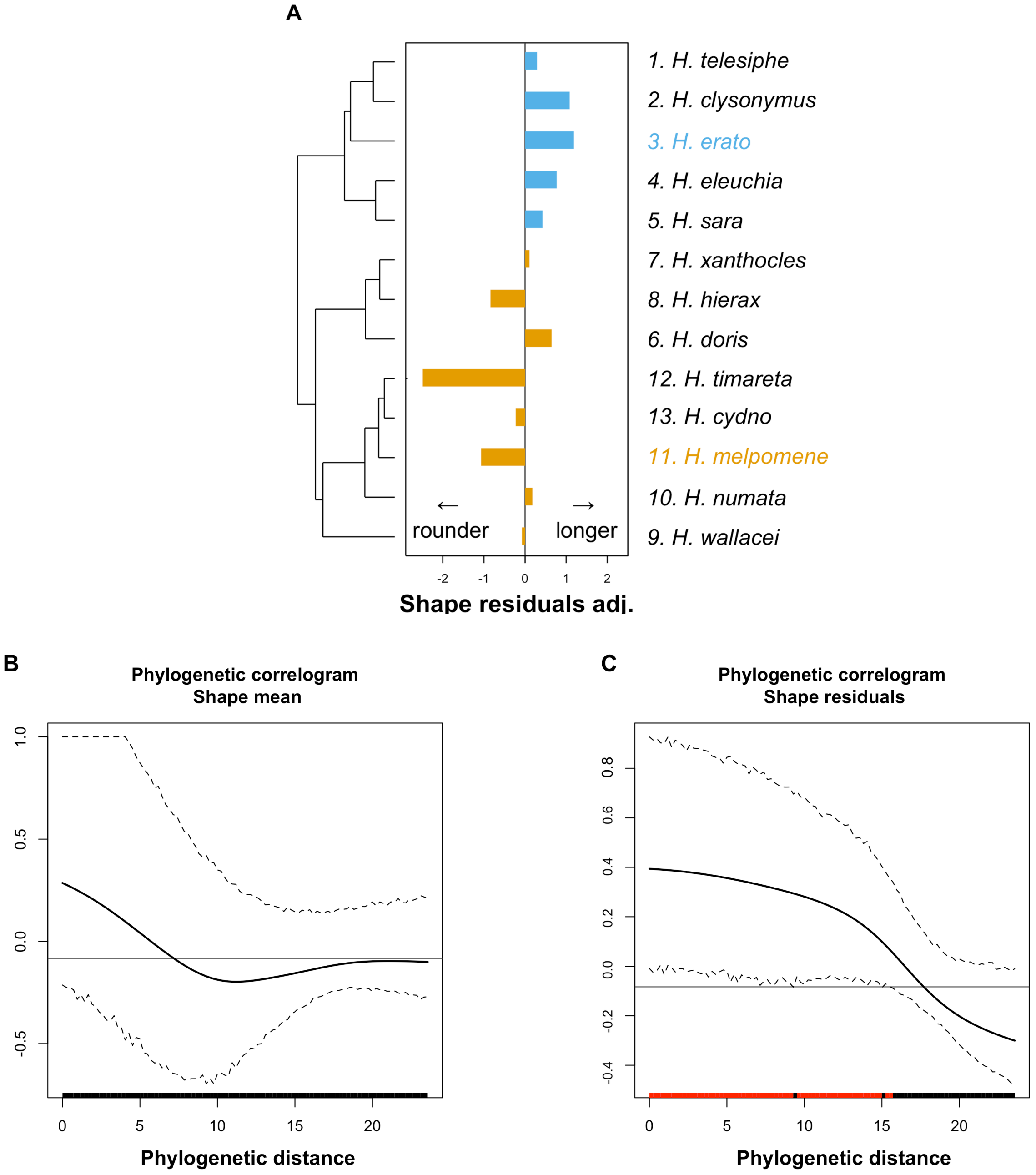


**Figure S2.** Phylogenetic signal in wing shape. A) Z-transformed wing shape residuals across the *Heliconius* phylogeny. B) phylogenetic correlogram of species mean wing shape. C) phylogenetic correlogram of species wing shape model residuals. The solid black line represents Moran’s I index of autocorrelation and the dashed black lines represent the lower and upper bounds of the confidence 95% confidence interval. The horizontal black line represents the expected value of Moran's I under the null hypothesis of no phylogenetic autocorrelation. The coloured bars in the x-axes show whether the autocorrelation is significant (based on the confidence interval): red for significant positive autocorrelation and black for nonsignificant autocorrelation. All figures were obtained with the package phylosignal **(Keck *et al.*, 2016).**

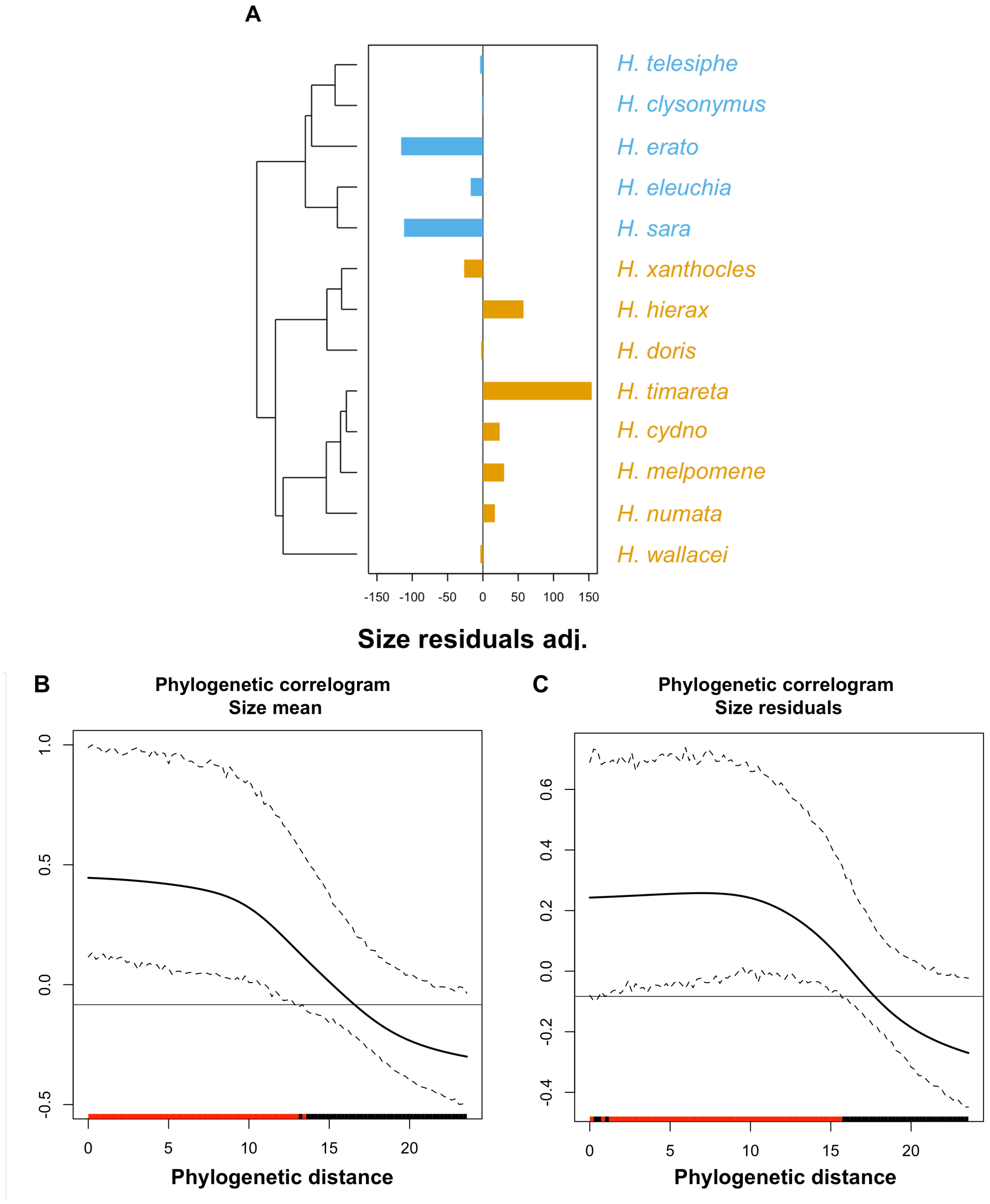


**Figure S3.** Phylogenetic signal in wing size. A) Centered wing size residuals across the *Heliconius* phylogeny. B) phylogenetic correologram of species mean wing size. C) phylogenetic correologram of species wing size model residuals. The solid black line represents Moran's I index of autocorrelation and the dashed black lines represent the lower and upper bounds of the confidence 95% confidence interval. The horizontal black line represents the expected value of Moran's I under the null hypothesis of no phylogenetic autocorrelation. The coloured bars in the x-axes show whether the autocorrelation is significant (based on the confidence interval): red for significant positive autocorrelation and black for nonsignificant autocorrelation.

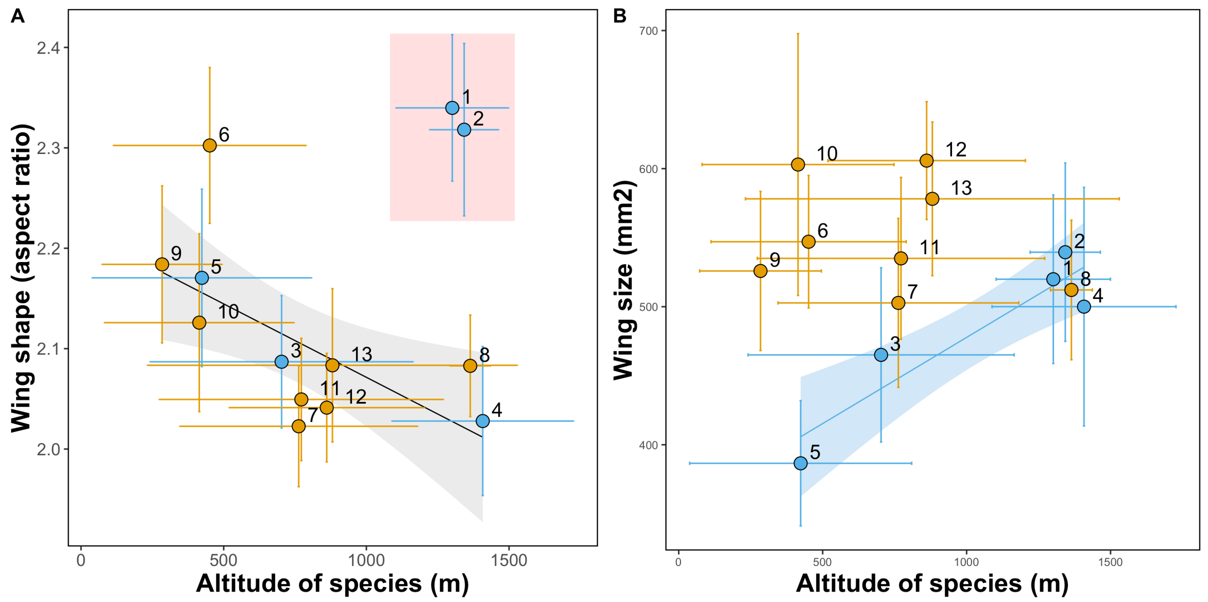


**Figure S4.** Species variation in raw wing shape (A, aspect ratio) and wing size (B, area in mm$^2$). Point represent species mean values. Horizontal and vertical lines show standard error for species' altitude and trait means, respectively. Lines show best linear fit for significant effects and are coloured by clade in B) (blue: erato clade, orange: melpomene clade). In A), fitted linear model (black line) excluded two species (1 and 2, in red panel) that had high Moran's I phylogenetic autocorrelation levels. Shaded areas show confidence bands at 1 standard error.  Numbers correspond to *Heliconius* species: 1 *H. telesiphe*, 2 *H. clysonymus*, 3 *H. erato*, 4 *H. eleuchia*, 5 *H. sara*, 6 *H. doris*, 7 *H. xanthocles*, 8 *H. hierax*, 9 *H. wallacei,* 10 *H. numata*, 11 *H. melpomene*, 12 *H. timareta*, 13 *H. cydno*.

**Table S2.** Phylogenetic Generalised Least Squares full model summaries for sexual size dimorphism, wing shape and wing size. Correlation structures of the models are shown in the third column. Dist. Eq.= distance from Equator, SD= sexual dimorphism.

| Response variable (wing trait) | Model type | Corr. structure | Fixed effects | Estimate | SE | t-value | p-value | d.f. (d.f. res.) |
|---|---|---|---|---|---|---|---|---|
| Size Sexual Dimorphism | PGLS (nmle) | Phylogeny, intra-sp variance, sample size | (Intercept) | 5.22 | 2.21 | 2.35 | 0.04 | 13 (10) |
| | | | Solitary larvae | -13.5 | 2.60 | -5.17 | 0.0003*** | |
| Shape | PGLS (nmle) | Phylogeny, intra-sp variance, sample size | (Intercept) | 1.25 | 0.32 | 3.89 | 0.006 | 13 (6) |
| | | | Size | 1.3E-3 | 4.4E-4 | 3.03 | 0.02* | |
| | | | Sex ratio | 0.88 | 0.175 | 5.03 | 0.002** | |
| | | | Dist. Eq | -3.6E-2 | 0.014 | -2.63 | 0.034* | |
| | | | Altitude | -2.4E-4 | 6.6E-5 | -3.62 | 0.008** | |
| | | | Adult-mating clade | -0.23 | 0.09 | -2.52 | 0.040* | |
| Size | PGLS (nmle) | Phylogeny, intra-sp variance, sample size | (Intercept) | -758 | 434.7 | -1.75 | 0.141 | 13 (6) |
| | | | Dist. Eq | 6.93 | 4.97 | 1.39 | 0.222 | |
| | | | Size SD | -24.3 | 11.2 | -2.17 | 0.081 | |
| | | | Sex ratio | 9.23 | 143.2 | 0.06 | 0.951 | |
| | | | Altitude | 1.34 | 0.51 | 2.63 | 0.047* | |
| | | | Shape | 584 | 229 | 2.54 | 0.052· | |
| | | | Altitude*Shape | -0.61 | 0.23 | -2.60 | 0.047* | |
| | | | Size SD*Sex ratio | 25.13 | 15.3 | 1.63 | 0.16 | |

#### Partial $R^2$ PGLS models

Phylogenetic signal gives a measure of the magnitude of the effect of shared ancestry, while partial $R^2$ of PGLS models can give information about the statistical significance of phylogenetic correlations on the model (Ives, 2018). Deriving partial $R^2$ can help tease apart how the predictor variables (fixed effects) interact with the phylogenetic correlation (random effects) in a PGLS model for the traits under study, even if total $R^2$ is not a useful goodness of fit metric for GLS models with correlated data, as model-fitting is highly stochastic (Ives, 2018).

**Table S3.** Partial R2 estimation for the different components of the PGLS model explaining natural variation in wing shapes across species. In the simplified models below, fixed effects included in the final model are represented by x, and the phylogenetic correlation is represented by phy.

| | Full model | Reduced model | $R^2$ |
|---|---|---|---|
| 1 | shape ~ x + phy | shape ~ 1 + phy | 0.79 |
| 2 | shape ~ x + phy | shape ~ x | $<0$ |
| 3 | shape ~ x | shape ~ 1 | 0.81 |
| 4 | shape ~ 1 + phy | shape ~ 1 | $<0$ |

What can be concluded from $R^2$ estimations in wing shape models (Table S3) is that removing the phylogenetic correlation from the full model would not decrease our explanatory power (row 2), and that phylogeny alone explains none of the variation in wing shape (row 4) whereas fixed effects alone explain 80% of the variation (row 3). Furthermore, the inclusion of fixed effects with a phylogenetic correlation structure adds much explanatory power (row 1) similarly to the inclusion of fixed effects alone (row 3). In this case, the effect of phylogeny on the fixed effects is not strong. Nevertheless, since the residuals of the wing shape GLS accounting for fixed effects had a strong phylogenetic signal (Fig. S2), we deemed necessary to retain the phylogenetic term.

**Table S4.** Partial $R^2$ estimation for the different components of the PGLS model explaining natural variation in wing sizes across species. In the simplified models below, fixed effects included in the final model are represented by *x*, and the phylogenetic correlation is represented by *phy*.

| | Full model | Reduced model | $R^2$ |
|---|---|---|---|
| 1 | size ~ x + phy | size ~ 1 + phy | 0.86 |
| 2 | size ~ x + phy | size ~ x | $<0$ |
| 3 | size ~ x | size ~ 1 | 0.93 |
| 4 | size ~ 1 + phy | size ~ 1 | 0.27 |

Wing size alone was shown to have a strong phylogenetic signal, so it is expected for the phylogenetic structure alone to explain some of the variation in size, in this case 31% (Table S4, row 4). Nevertheless, removing the phylogenetic correlation does not reduce the explanatory power of the model when fixed effects are being accounted for (row 2), whereas removing the fixed effects reduces the explanatory power by 84% (row 1). Nevertheless, since the residuals of the wing size GLS accounting for fixed effects had a strong phylogenetic signal (Fig. S2), we deemed necessary to retain the phylogenetic term.
